# IRescue: uncertainty-aware quantification of transposable elements expression at single cell level

**DOI:** 10.1101/2022.09.16.508229

**Authors:** Polimeni Benedetto, Marasca Federica, Ranzani Valeria, Bodega Beatrice

**Affiliations:** INGM, Istituto Nazionale di Genetica Molecolare ‘Romeo ed Enrica Invernizzi’, Milan, Italy; Ph.D. Program in Translational and Molecular Medicine, DIMET, University of Milan-Bicocca, Monza, Italy; Department of Clinical Sciences and Community Health, University of Milan, Milan, Italy; Department of Biosciences, University of Milan, Milan, Italy

## Abstract

Transposable elements (TEs) are mobile DNA repeats that contribute to the evolution of eukaryotic genomes. In complex organisms, TE expression is tissue specific. However, their contribution to cellular heterogeneity is still unknown and challenging to investigate in single-cell RNA sequencing (scRNA-seq), due to the ubiquity and homology of TEs in the genome. We introduce IRescue (Interspersed Repeats single-cell quantifier), the first software that accurately estimates the expression of TE subfamilies at single-cell level, implementing a UMI deduplication algorithm to allocate reads ambiguously mapped on TEs, while correcting for UMI sequencing errors. Applying IRescue on simulated datasets and real scRNA-seq of colorectal cancers, we could precisely estimate TE subfamilies expression. We show that IRescue improves the definition of cellular heterogeneity, detecting TE expression signatures and specific TE-containing splicing isoforms.

## Introduction

Transposable elements (TEs) are mobile genetic elements found in the genome of most eukaryotes. The genomic prevalence of TEs largely varies between organisms (1), and make approximately 46% of the human genome (2). In most species, retrotransposons are by far the most abundant TEs, as they can mobilize and amplify themselves thanks to a “copy-and-paste” replication mechanism (3). Based on their insertion age, TEs have been hierarchically organized into classes (e.g. LINE, SINE, LTR), superfamilies (e.g. LINE1, Alu, ERVL) and families or subfamilies (e.g. L1PA2, AluY, HERVL) (4). The LINE1 superfamily comprehends the last few autonomously active and mobile TEs in the human genome (5, 6). Indeed, the vast majority of TEs have lost the ability to generate new insertions; however, they can still be transcribed within surrounding transcriptional units and provide regulatory elements, affecting gene expression and RNA processing (7, 8). TEs are transcribed in a tissue specific fashion (9) and transcripts derived by TEs are involved in the epigenetic regulation of cell identity and differentiation (10, 11). However, the impact of TE expression in cellular heterogeneity at single cell level has yet to be investigated.

Next Generation Sequencing (NGS) technologies were indispensable to discover and annotate TE insertions in reference genomes and to realize the impact of TEs in genome and transcriptional regulatory networks. Yet, the study of TEs with NGS is hampered by their repetitive nature and high degree of homology between elements (12). Some precautions during library design, such as choosing a paired-end layout, increasing read length (13) and using specific software (14, 15), can drastically improve both read mappability and expression estimate of TEs. While a plethora of software are available for bulk RNA-Seq, few attempts have been made for measuring the expression of TEs in single-cell RNA sequencing (scRNA-seq) so far. The vast majority of scRNA-seq libraries in public repositories are derived from droplet-based technologies (e.g. most Chromium 10x, Drop-seq and inDrops kits) (16), and thus usually characterized by short reads with a strong 3’- or 5’-end positional bias. Moreover, these reads are effectively single-end, since the cDNA insert is only represented by one mate, while the other carries the cell barcode and unique molecule identifier (UMI) sequences. This tag-based layout decreases read mappability and makes it difficult to determine the exact genomic origin of TE-containing RNA fragments. Current tools for the quantification of TE subfamilies only use one alignment per UMI, disregarding the signal from ambiguous alignments on different TE subfamilies and do not take into account sequencing errors in UMI sequences (17, 18), that are common and introduce an overestimation of the UMI count (19). Approaches to rescue ambiguously mapping UMIs and obtain better expression estimates only exist for gene expression quantification, and leverage on finding UMI-transcript equivalence classes (20, 21). Here we present IRescue (interspersed repeats single-cell quantifier), a command-line tool for the error-correction, deduplication and quantification of UMIs mapping on TEs in scRNA-seq using a UMI-TE equivalence class-based algorithm. IRescue is currently the only software that, in case of UMIs mapping multiple times on different TE subfamilies, takes into account all mapped features to estimate the correct one, rather than excluding multi-mapping UMIs or picking one random alignment per UMI.

In this study, we demonstrate the precision of IRescue using simulated data and identify TE expression signatures associated with colorectal cancer (CRC) in real datasets. Moreover, we show that using IRescue it is possible to dissect the expression dynamics across distinct CRC cellular subsets of tumor-specific TEs previously identified in bulk RNA-seq data (22, 23). Finally, we suggest that the expression of these marker TEs is explained by the transcription of TE-containing tumor-specific alternative isoforms of human oncogenes.

## Materials and Methods

### IRescue workflow

#### Input data

IRescue requires as input a binary aligned map (BAM) file (24) containing read alignments on a reference genome, which must have the cell barcode and UMI sequences annotated as key-value tags. The keys for both the cell barcode and the UMI are by default the strings “CB” and “UR”, with the possibility to be overridden by the user. Additionally, it is mandatory to provide either the name of a genome assembly (e.g. “hg38” for the human genome) or a browser extensible data (BED) file (25) containing genomic TE coordinates. If a genome assembly name is provided, IRescue will automatically download and parse the Repeatmasker (26) coordinates from the UCSC servers at the beginning of the workflow, filtering out unwanted short tandem repeats and repetitive RNA classes (i.e. Low_complexity, Simple_repeat, rRNA, scRNA, srpRNA, tRNA). If a custom BED file is provided, it is advised to use the TE subfamily as the feature name in the fourth column. The BED file must at least have four columns, can be plain text or compressed and the chromosome names must follow the same nomenclature as the genome assembly used for alignment. Repeatmasker annotations automatically downloaded by IRescue follow the UCSC nomenclature, i.e. with reference names having the “chr” prefix. Optionally, a cell barcode whitelist file (either plain text or compressed) can be provided in order to filter the reads with valid cell barcodes only. We recommend mapping reads with a spliced aligner that reports the sequence-corrected cell barcodes, such as the “CB” tag added by CellRanger (27) or STARsolo (28, 29), and to provide the filtered barcode list (i.e. the “barcodes.tsv” file) as a whitelist to IRescue, in order to obtain a TE count matrix consistent with the gene count matrix provided by these tools (Supplementary Figure S1A).

#### Mapping alignments on TEs

Read alignments and TE coordinates are processed in parallel by chromosome, according to the number of allocated CPUs. The intersection between read and TE coordinates is performed by IRescue wrapping bedtools 2.30.0 (30), taking into account eventual read deletions and splitting events due to splice junctions (i.e. TE loci localized between donor and acceptor splicing coordinates are not considered mapped). In case of UMIs aligned on multiple TE loci, all the aligned TE features will be parsed and carried over for the UMI deduplication.

#### UMI deduplication algorithm

After mapping, cell barcodes are evenly splitted up in batches for parallel processing of single cell TE counts, according to the number of allocated CPUs. For each cell, UMIs and TE features are indexed and stored in memory as key-value pairs. UMIs mapped on the same set of TE features (i.e. keys having the same value) are stored in the same equivalence class (EC), with the assumption that UMIs mapping on the same set of interspersed repetitive features carry an “equivalent” information regarding the TE-derived RNA molecule of origin. For each EC, the sequences of UMIs are compared in a pairwise manner to calculate the number of mismatches between them. Next, UMIs are arranged in an undirected graph where each node is a UMI and edges connect UMIs that differ for just one mismatch, as they are considered potential duplicates. To calculate the deduplicated number of UMIs explaining the EC, the algorithm finds the neighborhood of each node, i.e. a set containing the node itself and its adjacent nodes. We discern three basic configurations of UMI networks. i) If a node has no adjacent nodes, thus a neighborhood of size (i.e. cardinality) one, it is considered a non-duplicated UMI, and adds 1 to the final UMI count of the EC. ii) If two or more nodes adjacent to each other all have the same neighborhood, they are considered derived from the same UMI, and adds 1 to the final UMI count of the EC. iii) In case of ambiguous connections, e.g. when three or more nodes that aren’t all adjacent are connected through a path, the algorithm finds the hubs of the graph, i.e. the nodes whose neighborhood’s intersection with any other neighborhood always results in a set of size smaller than the hub’s neighborhood’s size (Supplementary Figure S1B). Formally, given a set of *n* nodes, with *n* ≥ 3, a neighborhood *N*_1_ is a hub if the following comparison is always true:

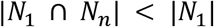

For example, in the graph in figure 1, the nodes 1 to 4 are connected on the same path, with nodes 1 and 3 being the network’s hubs, thus adding 2 to the UMI count, whereas the nodes 5 and 6 are a couple of adjacent nodes with the same neighborhood, hence derived from the same UMI.

**Figure 1.**
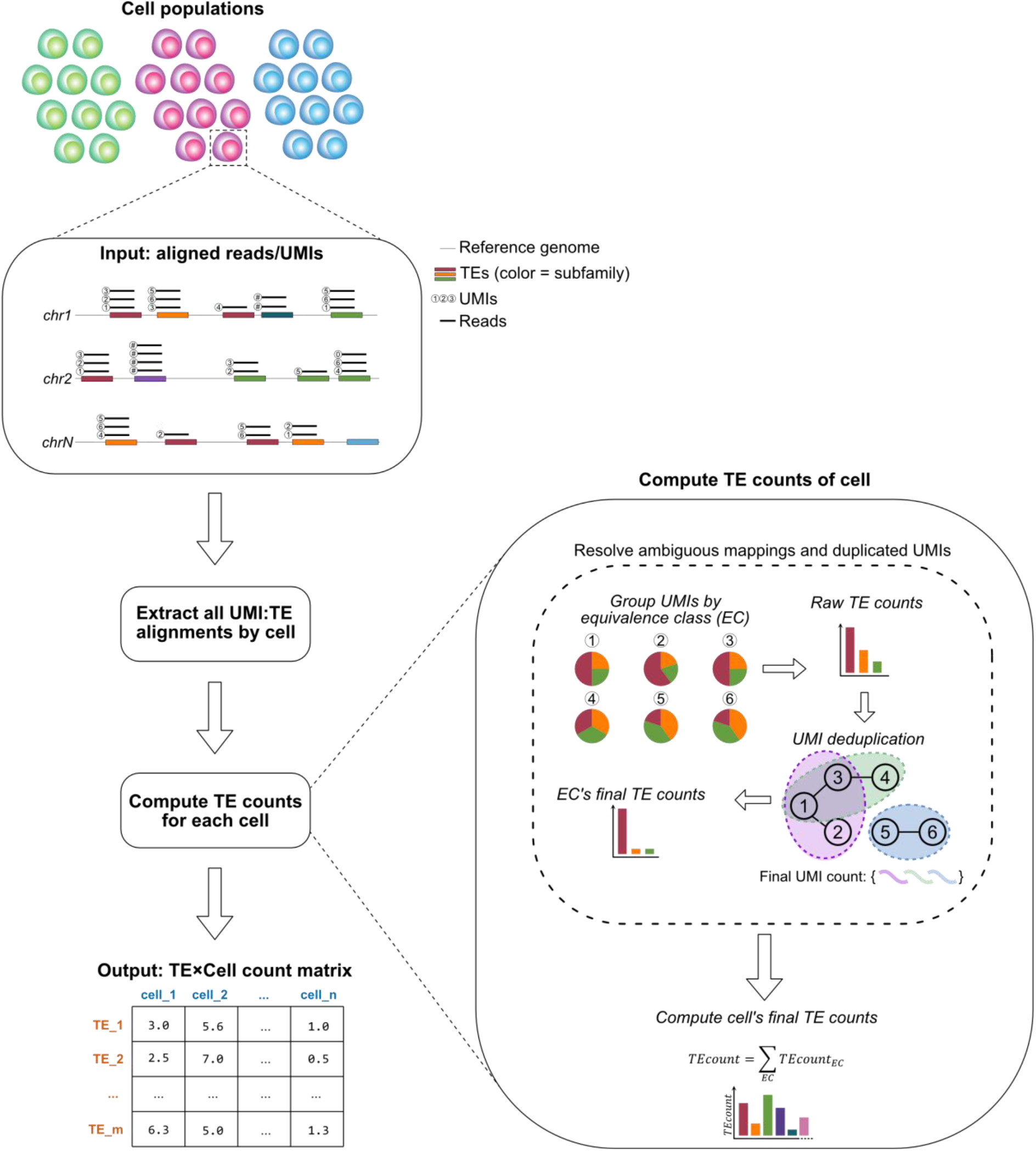
IRescue algorithm overview. scRNA-seq reads aligned on a reference genome, and the associated UMIs and cell barcodes, are parsed by IRescue and, for each cell, UMIs mapped on the same set of TEs are grouped into equivalence classes (ECs). For each EC, the final UMI count is inferred by deduplicating the UMIs based on their sequence similarity. The deduplicated UMI count is assigned on the TE feature showing the highest amount of alignments (in case of ties, the count is splitted between features). The counts of all ECs of a cell are summed together to obtain the cell’s final TE counts. The TE counts are written in a m×n matrix, with m and n being the number of TEs and cells, respectively.

#### TE features count

Once the EC’s UMI count has been obtained, it is assigned to the TE feature showing the highest number of alignment events in the EC. Finally, the count of TE features obtained from every EC of a cell is summed up, to obtain the final TE counts of the cell:

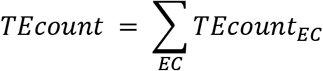

The counts of TE features across cells are written in a TE×Cell sparse matrix following the Market Exchange format (MEX), which is compliant with the gene matrix output of Cell Ranger or STARsolo to ensure compatibility with several toolkits for single cell downstream analysis.

### scRNA-seq alignment and gene quantification

Read alignments and single cell gene counts matrix were obtained by mapping scRNA-seq reads to the reference human genome (UCSC hg38 primary assembly) using STARsolo 2.7.9a (28, 29). As genes and splice junction database, we provided the Gencode comprehensive annotation v32 (31) with chromosome names converted to the UCSC nomenclature (passed to STAR through the `“––sjdbGTFfile”` parameter). We provided the 10x Genomics cell barcodes whitelist for the v2 library kit (passed through the `“––soioCBwhiteiist”` parameter). Other non-default parameters were: “––outSAMattributes NH HI AS nM NM MD jM jI XS MC ch cN CR CY UR UY GX GN CB UB sM sS sQ ––outFilterMultimapNmax 100 ––winAnchorMultimapNmax 100 ––twopassMode Basic ––soloType CB UMI Simple ––soloCellFilter EmptyDrops_CR 10000 0.99 10 45000 90000 500 0.01 20000 0.01 10000”.

### Single cell TE expression quantification

To estimate the expression of TEs at single cell level, read alignments were processed using IRescue, providing the filtered barcodes list produced by STARsolo as cell barcodes whitelist (passed through the `“––whitelist”` parameter), and other parameters `“––genome hg38 ––CBtag CB ––UMItag UR ––keeptmp”`. To compute TE counts with scTE 1.0 (17), the same Repeatmasker and Gencode annotations were used to build an index, keeping other parameters as default, and read alignments were counted with parameters `“–CB CB–UMI UR”`.

### Data simulations and benchmarks

Simulated scRNA-seq reads and TE subfamily counts were obtained adapting the method described by Kaminow et al. (28), which allows to reproduce the positioning bias of tag-based scRNA-seq on genes, introns and intergenic regions by using a real dataset as the basis for the simulation. For this purpose, we used a 10x Genomics v2 3’-end PBMC dataset (https://www.10xgenomics.com/resources/datasets/8-k-pbm-cs-from-a-healthy-donor-2-standard-2-1-0) (32), which is a common scRNA-seq library layout. Briefly, cell barcodes are filtered according to the 10x Genomics whitelist and UMIs with uncalled bases are removed, reads are aligned on a reference that combined the human hg38 primary genome assembly and the Repeatmasker TE genomic sequences using BWA-MEM 0.7.17 (33). UMIs are counted based on the TE subfamily they map on; in case of alignments on multiple features, the top-scoring alignment is chosen. Finally, the aligned reference sequences were extracted and a mismatch error rate of 0.5% was added to simulate Illumina sequencing errors. Plots summarizing multimapping UMIs and equivalence classes were obtained from the UMI-TE mappings file generated by IRescue using the `“––keeptmp”` parameter. Spearman correlation tests were done using R 4.1.0 (34) between IRescue and simulated TE counts or features per cell, or between IRescue and scTE total counts or counts by cell. The relative deviation (RD) between measured and simulated counts was calculated as:

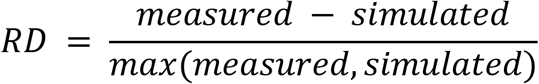

and represented on a histogram with 0.001 bin size. Cell clustering according to TE counts was performed using the Louvain algorithm implemented in Seurat 4.0.5 (35). The comparison between IRescue or scTE and simulated clusters was performed with a chi squared test, and visualized by plotting the Pearson’s residuals using the R package corrplot 0.92 (36). Memory (RAM) usage and run time were measured by running IRescue and scTE in a Nextflow pipeline (37) and extracted from the pipeline’s report.

### Analysis of scRNA-seq cancer dataset

Cancer and normal scRNA-seq raw data were obtained from ArrayExpress E-MTAB-8410 (38), along with the annotation of cell identity based on gene expression. The epithelial cells from six paired tumor and normal samples were processed with IRescue for quantifying TE expression. TE counts normalization, Principal Component Analysis (PCA), Cell clustering (using the Louvain algorithm) and Uniform Manifold Approximation and Projection (UMAP) were performed using the Seurat 4.0.5 toolkit. Cell clusters based on TE expression profiling were annotated as tumoral (K) or normal (N) according to the most prevalent cell condition on each cluster. TE expression signatures were obtained by finding differentially expressed TEs between each cluster and the rest of the dataset, and TEs significatively enriched (Wilcoxon rank sum test’s p-value adjusted according to Bonferroni < 0.05; log2 fold change > 1) on K or N clusters only were selected, discarding TEs enriched in both conditions. The difference in enrichment of TE subfamilies by class between tumor and normal signatures was tested with a two-tailed two-proportions Z-test using the R 4.1.0 stat package. The average expression of CRC marker TEs across conditions or clusters was visualized using Seurat 4.0.5. Sashimi plots were obtained with the IGV genome browser (39).

## Results

### IRescue: an algorithm for TE expression quantification using ambiguously mapped UMIs in scRNA-seq

Rescuing multimapping reads is well known to be a compulsory procedure in order to accurately quantify the expression of TEs (40) or other poorly mappable features, such as multigene families (41). Hence, one of the cornerstones that motivated the development of IRescue was to use the information of all mapped features to allocate ambiguously mapped UMIs with higher confidence. IRescue allows integrating TE and gene counts in a canonical scRNA-seq workflow analysis (Supplementary Figure S1A) and contains a novel UMI deduplication algorithm that takes into account mismatches between both uniquely and multimapping UMIs. IRescue takes as input reads aligned on a reference genome assembly with cell barcode and UMI annotated as tags of type string, and genomic TEs coordinates. While any spliced aligner generating an output with these features will be compatible with IRescue, as of today we recommend using STARsolo (28, 29) with custom parameters optimized for reads alignment on interspersed repetitive elements (see methods). IRescue is programmed to download and parse into a BED file the genomic Repeatmasker TE coordinates (26) of the genome assembly of choice directly from the UCSC server (25); alternatively, it is possible to use a custom BED file with TE coordinates. The output of IRescue is a sparse matrix written in a Market Exchange Format file (MEX), compliant with the 10x Genomics Cell Ranger’s output (27) to ensure compatibility with most toolkits for downstream analysis (35, 42) (Supplementary Figure S1A). IRescue extracts all the alignments on TEs and parses the cell barcode and UMI sequences, as well as the mapped TE features; then, computes the TE counts for each cell by running an algorithm for the correction of multimapping reads and duplicated UMIs. Briefly, UMIs mapped within a cell on the same set of TE features are grouped into *n* equivalence classes (ECs); then, for each EC of a cell, duplicated UMIs are detected by inspecting their sequence similarity, admitting up to one mismatch in UMI sequence to find duplicates. After UMI deduplication, the final UMI count for an EC is assigned to the TE feature showing the highest number of alignments (in case of ties, the count gets evenly splitted among the features). The counts of TE features from all ECs within a cell are summed up to obtain the cell’s final TE counts. Finally, the TE counts of all the sample’s cells are collected and written into a TE×Cell matrix (Figure 1). To limit the TE expression quantification to valid cell barcodes, it is possible to give as an additional input to IRescue a cell barcode whitelist obtained using the whitelisting method implemented by the gene expression quantifier of choice (27, 28, 43–45). In this paper, we provided as a whitelist the filtered barcodes list obtained from STARsolo during the alignment pre-processing step using the EmptyDrops algorithm (45). IRescue is written in the Python programming language and leverages on essential open source libraries for efficient scientific computing and bioinformatics data wrangling (24, 30, 46–48).

### scRNA-seq simulations confirm that IRescue is accurate in quantifying TE expression

To evaluate the precision of IRescue’s quantification strategy, we sought to generate simulated scRNA-seq reads carrying the same biases of real datasets, such as reads mapping to multiple genomic loci and to intronic and intergenic regions, including TE fragments spliced in unknown transcript isoforms or autonomously transcribed TEs. To achieve this, we implemented the method described by Kaminow et al. (28), in which they used a real dataset as a basis for the generation of simulated reads and counts, with custom modifications in order to obtain simulated counts at TE subfamily level, rather than gene-level (see methods). As the basis for the simulation, we used a typical 10x Genomics v2 protocol library of peripheral blood mononuclear cells, containing 8,341 cells with ~92,000 reads per cell, 16bp long cell barcodes, 10bp UMIs and 98bp 3’-end cDNA sequences (32). After aligning simulated reads on the reference genome, we assessed the mappability over TEs at UMI-level and found that, of the 33% UMIs mapped on TEs, only 21% were uniquely mapped, while 39%mapped multiple times on the same TE subfamily and the remaining 40% on different TE subfamilies (Figure 2A). We obtained TE counts from the aligned simulated reads using IRescue. To evaluate the distribution and composition of ECs across cells, we processed IRescue’s intermediate output and observed that most cells contains ~1000 ECs, each consisting of a unique set of UMIs mapping on the same set of TE subfamilies (Figure 2B), and that most ECs contains 10 or less different TE subfamilies (Figure 2C). Therefore, UMIs mapping on more than 10 distinct TE subfamilies are rare. To test the accuracy of estimating TE expression in single cells, we calculated the correlation between simulated and measured TE counts (Figure 2D) and number of TE subfamilies (Figure 2E) and found it very high in both measurements. We then compared the analysis on simulated data with scTE (17), and found a stronger correlation between measured and simulated counts using IRescue, compared to scTE (Supplementary Figure S2A), indicating a better accuracy for IRescue. Furthermore, TE counts measured across all cells showed a minimal deviation from the simulated counts in both IRescue and scTE (Supplementary Figure S2B), whereas cell’s total TE counts correlated better to simulated counts in more cells using IRescue, compared to scTE (Supplementary Figure S2C). To test the ability to infer the identity of cells in respect to their TE expression profile, we evaluated the performance of dimensionality reduction and cell clustering techniques on measured and simulated data. In order to group cells according to their TE expression profile, we applied a canonical workflow of scRNA-seq analysis using the Seurat toolkit (35, 49). We found that clusters of cells analysed with IRescue are equivalent to their simulated counterpart, with only a partial overlap observed between clusters 0 and 2 (Figure 2F). Performing the same analysis with scTE, we found that two of nine identified clusters completely failed to match a correspondent identity in simulated data, showing that IRescue performs better in identifying the cell’s state (Supplementary Figure S2D). Finally, we analysed the resource usage of IRescue and scTE by the number of processing units (CPUs) utilized. While scTE had a shorter run time when using a low number of CPUs, IRescue’s run times scaled better with an increasing number of CPUs (Supplementary Figure S2E); on the other hand, IRescue showed a very low memory footprint compared to scTE (Supplementary Figure S2F). The higher run time on IRescue is expected, as it performs more tasks than scTE (including UMI mismatch correction and weighting of multi-mapped TEs). Because IRescue is programmed to evenly split up the cells according to the allocated CPUs for parallelizing the count process, the run time drastically decreases when using a higher number of CPUs. The low memory footprint of IRescue is achieved by an efficient use of Python data structures for the indexing of cell barcodes, UMIs and TE features, and by performing operations leveraging on Python generators with a first-in-first-out logic.

**Figure 2.**
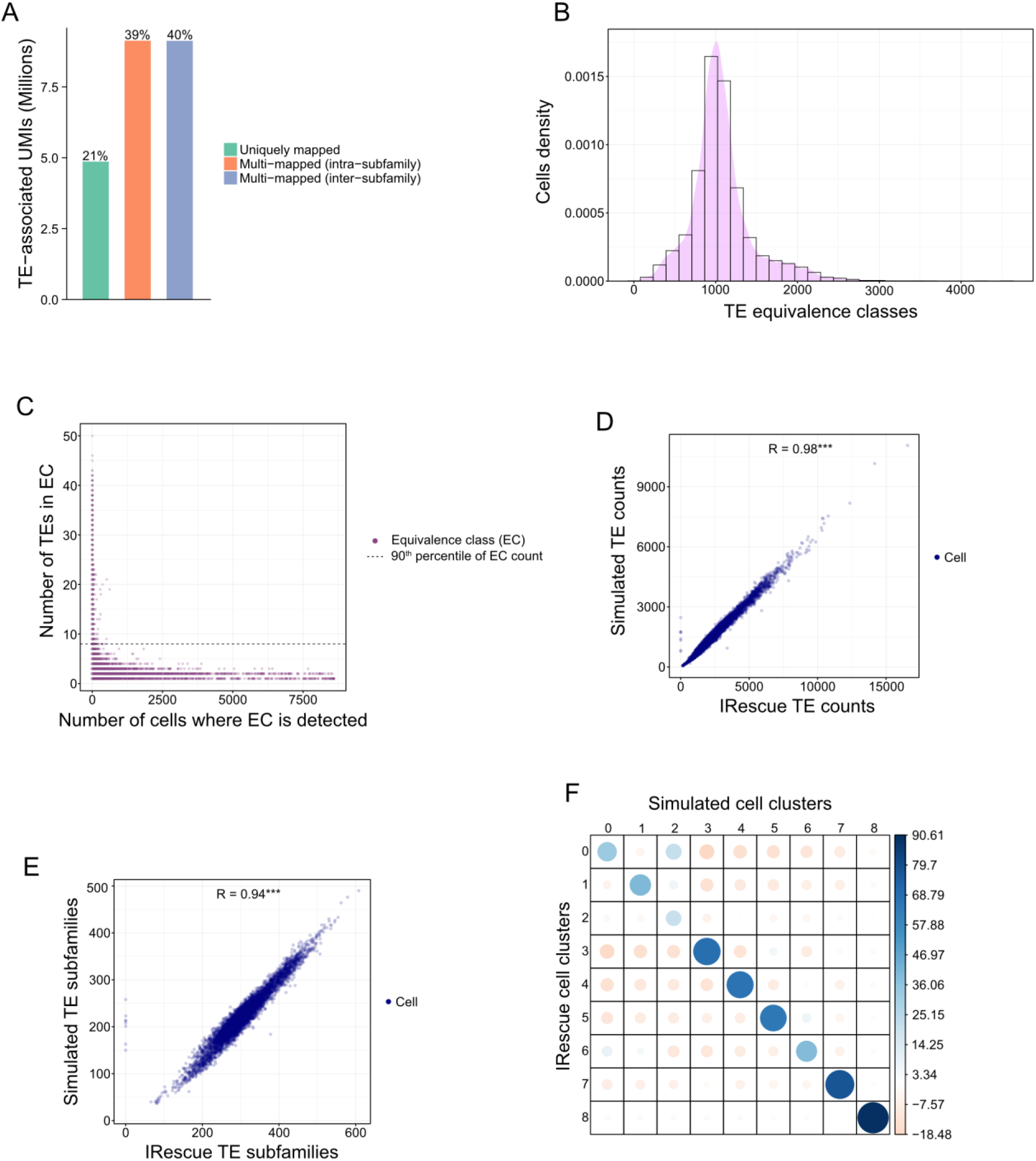
Performance evaluation of IRescue using simulated scRNA-seq data. (**A**) Count of TE-associated UMIs, divided into uniquely mapped (i.e. mapped only once in the genome), multi-mapped across multiple loci of the same TE subfamily (intra-subfamily) or across different TE subfamilies (inter-subfamily). (**B**) Distribution of the number of detected equivalence classes per cell. (**C**) Distribution of equivalence classes according to the number of cells on which they are detected and the number of TEs they contain. (**D-E**) Relationship between IRescue and simulated TE counts (D) or TE subfamilies (E). R = Spearman correlation coefficient. *** p-value < 2.2×10-16 (Spearman’s rank correlation test). (**F**) Correlation matrix plot between cell clusters inferred from IRescue and simulated TE expression values, using the Louvain algorithm. Pearson’s correlation residuals of all pairwise combinations are plotted as dots shaded blue or red depending on the correlation being positive or negative. Color intensity and dot size are proportional to the magnitude of the correlation. p-value < 2.2×10-16 (chi-squared contingency table test).

### IRescue enables the identification of tumor-specific TE signatures in colorectal cancer at single-cell resolution

TEs are known to be dysregulated in cancer, where they can initiate oncogene expression (50) or cause genomic instability (51, 52). Several studies reported the overexpression of specific TE subfamilies in cancer cells with bulk-level NGS analyses (22, 23, 50); however, how TEs are expressed in cancer at single cell level and whether they contribute to the heterogeneity of cell subsets is still poorly characterized. Therefore, to demonstrate IRescue performance in real datasets, we analysed publicly available 10x Genomics 3’ scRNA-seq datasets from tumor and normal adjacent tissues of six colorectal cancer (CRC) patients (38). We performed an unsupervised clustering analysis on 32,276 tumor and normal epithelial cells based on TE counts computed by IRescue. Interestingly, we found that cell clustering based on TE expression infers 12 clusters (Figure 3A), of which half are populated exclusively or mostly by cancer cells (K clusters) and the other half by normal cells (N clusters, Figure 3B), showing that TE expression alone discerns cancer from normal cells. To identify the TEs specifically expressed in cancer or normal condition, we obtained the TE expression signatures of tumor and normal cells clusters by performing a differential expression analysis and selecting the TEs overexpressed only in one of the two conditions. We found that expression of specific TE subfamilies is detectable in a larger fraction of cells belonging to cancer clusters, compared to the normal ones (Supplementary Figure S3A-B). Interestingly, we grouped the differentially expressed TE subfamilies by class and found that LINE subfamilies are significantly over-represented in the TE expression signature of cancer cells, compared to the normal one (Figure 3C), while the other TE classes are similarly distributed between conditions. By inspecting the differentially expressed LINE subfamilies, we observed that they all belong to the LINE1 superfamily; moreover, the most evolutionarily young ones (i.e. L1HS and L1PA*) are those overexpressed in cancer cells only (Figure 3D). Bulk RNA-seq studies have identified 16 TE subfamilies known to be overexpressed in CRC (22, 23). We checked the expression of these TEs across the cell clusters identified in scRNA-seq, and confirmed the presence of 10 out of 16 CRC marker TEs among the TE subfamilies belonging to the cancer TE signature (Figure 3E). Furthermore, thanks to the single-cell resolution, we were able to reveal a previously uncharacterized heterogeneous distribution of CRC markers TEs (Figure 3F). For instance, L1PA2 and HERVH-int, both known to be associated with CRC (23), are enriched in different clusters of cancer cells (Supplementary Figure S3C).

**Figure 3.**
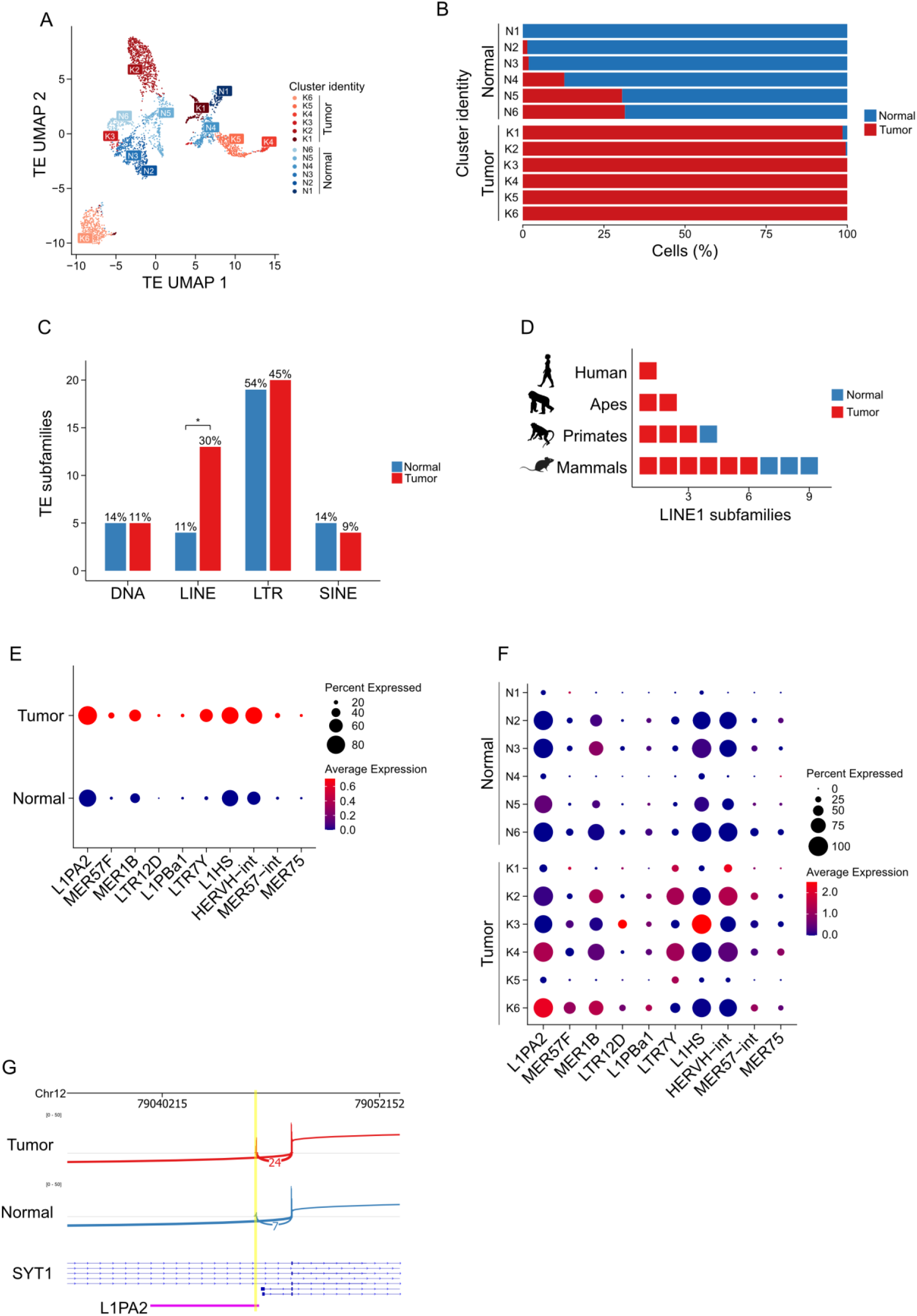
Identification of TE expression dynamics in colorectal cancer. (**A**) UMAP representation of CRC and normal cells according to TE expression. Clusters of normal and tumour cells (indicated in legend) are obtained on the basis of TE expression. (**B**) Barplot of the relative abundance of cells by condition across TE clusters. (**C**) Barplot of abundance (y-axis) and enrichment (percentage on top of bars) of TE subfamilies by TE class in tumor and normal TE signatures. * p-value < 0.05 (two-sided two-proportions Z-test). (**D**) Representation of LINE1 subfamilies specific for cancer or normal cells, stratified by the phylogenetic clade in which the insertions originated. (**E**) Dot plot of the average expression of known CRC markers TEs in tumor and normal conditions. The dot size is indicative of the percentage of expressing cells. (**F**) Dot plot of the average expression of known TE CRC markers across TE clusters. The dot size is indicative of the percentage of expressing cells. (**G**) Sashimi plot representing the coverage of a L1PA2-derived CRC-specific cryptic exon of SYT1 oncogene and the splitted reads across the cryptic and canonical exon.

Next, we asked whether it was possible to infer the structure of specific TE-containing transcripts. Assembling the entire structure of unannotated transcripts is unfeasible in droplet-based scRNA-seq, but it is still possible to inspect the usage of splice junctions (28). Since a specific L1PA2 insertion was reported to be spliced into an alternative isoform of the oncogene SYT1 in CRC (50), we inspected the read alignments over this particular locus, specifically looking for reads splitted across a splice junction mapped on both TEs and non-repetitive genomic regions that could support the existence of cryptic TE-containing exons. We found over 3-fold more reads splitted between the L1PA2 exon and the 3rd exon of SYT1 in tumor (Figure 3G), supporting the detection of the same previously reported L1PA2-derived tumor-specific cryptic exon (50). Likewise, we confirm the detection of a MER1B-PIWIL1 TE-oncogene isoform reported in CRC (50) (Supplementary Figure S3D). These results describe the complex dynamics of TE expression in CRC, that involve specific TE subfamilies across groups of cells, with a resolution that was not possible to achieve by previous works based on bulk-level NGS strategies. This supports the possibility to use TE subfamilies as markers to assist the dissection of cell heterogeneity. Moreover, these findings support the notion that the differential expression of TE subfamilies in cancer cells results from tumor-specific alternative splicing events of TE-containing transcripts.

## Discussion

In this work, we developed IRescue, the first software for error-correction, deduplication and quantification of UMIs mapping on multiple interspersed genomic features, such as those derived from transposable elements. It is known that reads originating from repetitive elements align equally well on multiple genomic loci (40), and aligners usually report either one random alignment per read, or one arbitrarily designated primary alignment and a number of secondary alignments. Therefore, discarding secondary alignments could lead to an incorrect feature assignment for that read. Several tools for TE expression quantification in bulk RNA-seq have been developed to take into account secondary alignments when assigning multi-mapping reads to features (14, 40). Such strategies have not yet been implemented in scRNA-seq, where only few methods for gene expression quantification are able to recover multi-mapping reads (28, 43, 44). In this context, IRescue not only takes into account all the secondary alignments of multi-mapping reads for feature assignment, but also provides the first strategy for the deduplication of UMIs multi-mapping on a reference genome. As a solution to assign a unique feature to multi-mapping UMIs, we leveraged on equivalence classes (ECs), a well-known concept used to count reads compatible to multiple isoforms in bulk RNA-seq (20, 21, 53), recently applied to gene expression quantification in scRNA-seq as well (43, 44).

We generated simulated 10x Genomics reads and TE subfamily counts and showed that, despite the high amount of ambiguous alignments, IRescue is highly precise when quantifying TE expression at subfamily level. We showed that, in a common 10x Genomics scRNA-seq library layout, one third of UMIs are TE-associated, of which 40% aligns on multiple TE subfamilies. This observation indicates that alignments on multiple TE meta-features are frequent and should be considered in TE expression quantification, even when performing a count at subfamily level. We compared IRescue to scTE (17), finding that, while TE counts are overall similar to simulations for both tools, IRescue performs better when inferring the cell’s identity using a common scRNA-seq clustering algorithm. We deem that this result is a consequence of scTE discarding the information of UMIs mapping to multiple TE subfamilies, an event that we showed being frequent in tag-based scRNA-seq.

In the context of TE overexpression in cancer, we analysed the single cell TE expression profiles of CRC and the tumor-adjacent tissue, and found an enrichment in evolutionarily young LINE subfamilies in the CRC TE signature. Furthermore, we showed that known CRC markers TEs, previously characterized in bulk RNA-seq, are heterogeneously expressed in different CRC cell clusters, and that the expression of such markers can be the result of the transcription of tumor-specific TE-containing alternative isoforms of human oncogenes (50). The transcription of RNA containing young LINE insertions in healthy somatic cells have been shown to be preferentially repressed during RNA processing, in contrast to old insertions that have been slowly co-opted by species throughout evolution (8). The transcription of TEs has been shown to be involved in several physiological scenarios, such as embryo development (11), aging (54), differentiation and cell identity (10), as well as pathological conditions, such as cancer (55) and neurodegenerative diseases (56); however, the expression dynamics and heterogeneity of the single cell TE transcriptome in most of these tissues is still unexplored. With the release of IRescue, we long to facilitate the single cell TE expression profiling from canonical scRNA-seq experiments, enabling the possibility of extracting novel information from the high amount of publicly available datasets (16).

## Data Availability

IRescue source code and documentation are available at https://github.com/bodegalab/irescue. Data used in this paper are available at 10x Genomics (https://www.10xgenomics.com/resources/datasets/8-k-pbm-cs-from-a-healthy-donor-2-standard-2-1-0) and ArrayExpress (E-MTAB-8410). The code to reproduce the results depicted in this paper is available at https://github.com/bodegalab/irescue_paper_analysis.

---

**Supplementary Figure 1.**
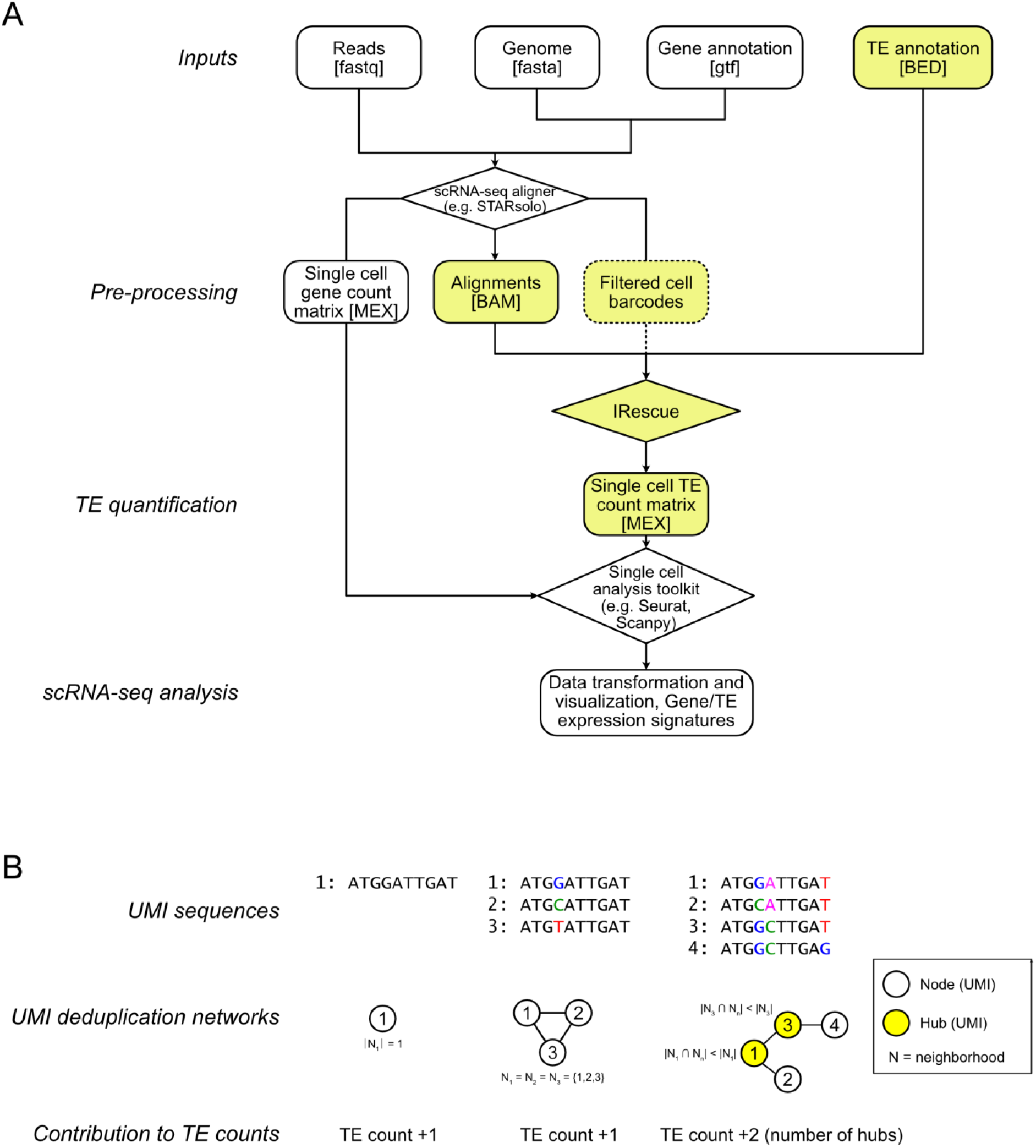
Integration of IRescue in a scRNA-seq pre-processing and analysis workflow. (**A**) Diagram depicting a workflow for scRNA-seq data analysis of TE expression using IRescue, where boxes represent data flow (with the data format indicated in squared brackets), diamonds represent the software used and highlighted boxes represent the portion of the workflow concerning IRescue. (**B**) Three exemplary UMI deduplication networks, where the contribution of the count of the equivalence class (EC count) is indicated at the bottom. From left to right: a UMI with no duplicates generates a graph of one node only; three duplicate UMIs with one-mismatch distance generate a graph with three nodes connected to each other; four UMIs with up to two-mismatch distance generate a complex graph whose contribution to the TE count is solved by identifying the graph’s hubs (yellow nodes).

**Supplementary Figure 2.**
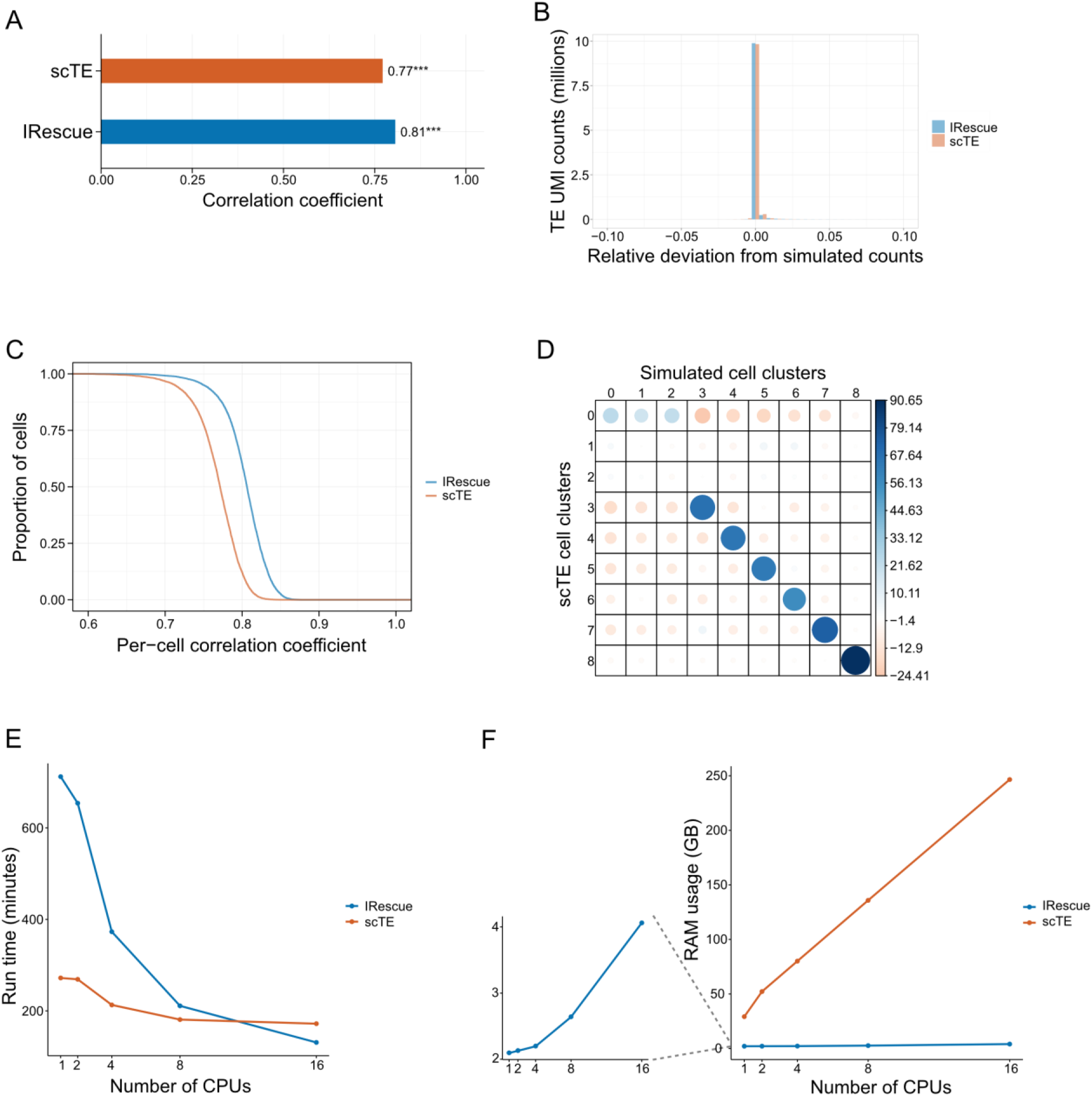
Accuracy assessments and benchmarking. (**A**) Spearman correlation coefficients of TE counts measured with IRescue or scTE against simulated counts (*** = Spearman’s rank correlation test’s p-value < 2.2×10-16). (**B**) Distance of TE counts measured with IRescue or scTE from simulated counts, expressed as relative deviation (see methods). Negative and positive values represent under- and over-estimated counts, respectively. (**C**) Reverse cumulative distribution of cells according to the Spearman correlation coefficient calculated between simulated and measured TE counts per cell. (**D**) Correlation matrix plot between cell clusters inferred from scTE and simulated data, using the Louvain algorithm implemented in Seurat v4. Pearson’s correlation residuals are plotted. Each pairwise combination is shaded blue or red depending on the correlation having a positive or negative sign, respectively, and the color intensity and size of the circles are proportional to the magnitude of the correlation. p-value < 2.2×10-16 (chi-squared contingency table test). (**E**) Run time of IRescue and scTE as a function of number of CPUs. (**F**) Memory usage of IRescue and scTE as a function of number of CPUs.

**Supplementary Figure S3.**
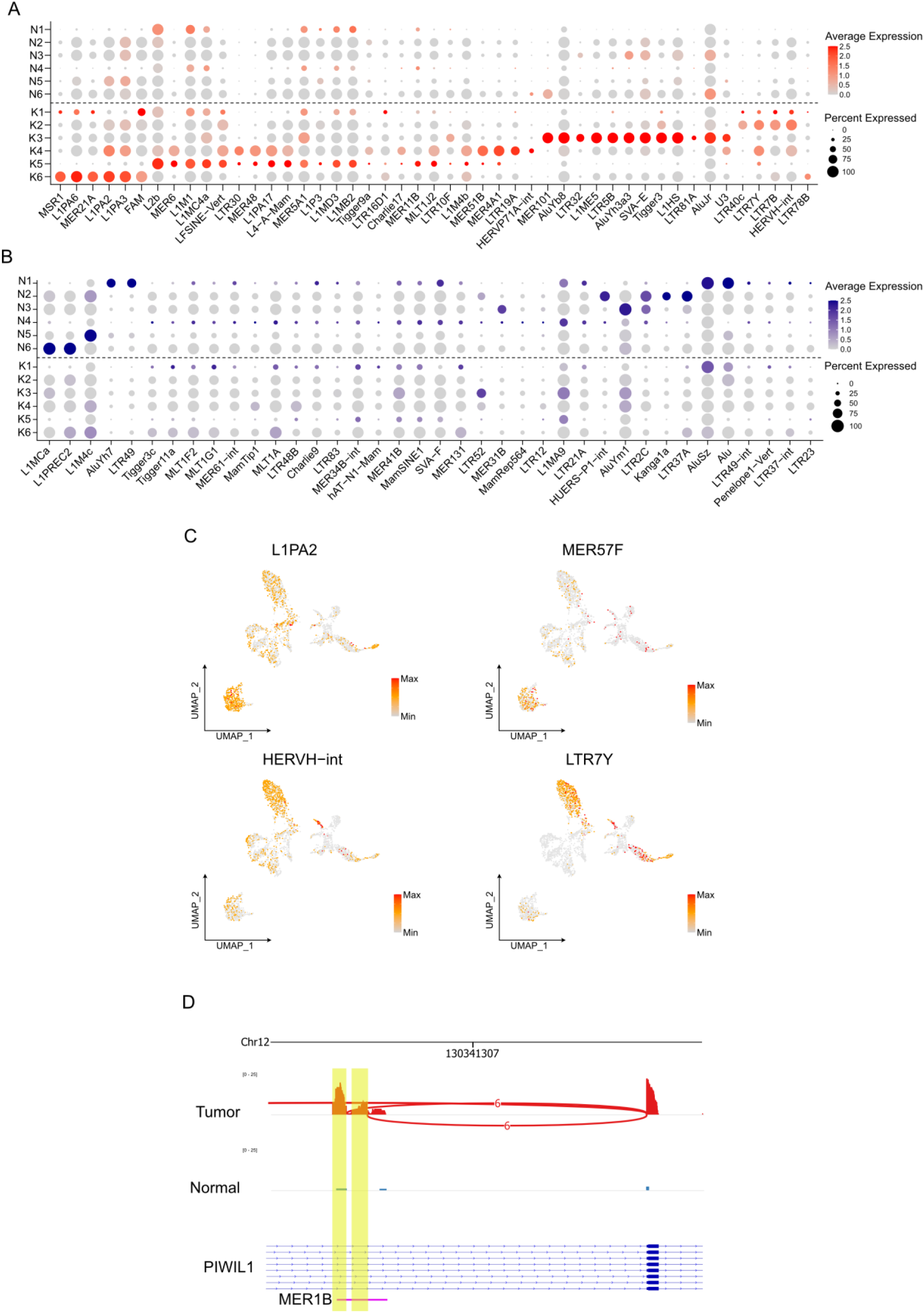
Identification of CRC cluster-specific TE subfamilies. (**A-B**) Dot plot representing the average expression of differentially expressed TEs in tumor (A) or normal (B) TE expression clusters (log2 fold change > 1 and adjusted p-value < 0.05, Wilcoxon rank sum test). The dot size is proportional to the percentage of expressing cells. (**C**) UMAP representation of CRC and normal cells according to TE expression. Color scale represent the scaled expression of representative CRC marker TEs. (**D**) Sashimi plot representing the coverage of a MER1B-derived CRC-specific cryptic exon of PIWIL1 oncogene and the reads splitted between the cryptic and canonical exons.

## References

1. Wells,J.N. and Feschotte,C. (2020) A Field Guide to Eukaryotic Transposable Elements. Annu Rev Genet, 54, 539–561.

2. Lander,E.S., Linton,L.M., Birren,B., Nusbaum,C., Zody,M.C., Baldwin,J., Devon,K., Dewar,K., Doyle,M., FitzHugh,W., et al. (2001) Initial sequencing and analysis of the human genome. Nature, 409, 860–921.

3. Kazazian,H.H. (2004) Mobile Elements: Drivers of Genome Evolution. Science, 303, 1626–1632.

4. Bao,W., Kojima,K.K. and Kohany,O. (2015) Repbase Update, a database of repetitive elements in eukaryotic genomes. Mobile DNA, 6, 11.

5. Brouha,B., Schustak,J., Badge,R.M., Lutz-Prigge,S., Farley,A.H., Moran,J.V. and Kazazian,H.H. (2003) Hot L1s account for the bulk of retrotransposition in the human population. Proc Natl Acad Sci U S A, 100, 5280–5285.

6. Beck,C.R., Collier,P., Macfarlane,C., Malig,M., Kidd,J.M., Eichler,E.E., Badge,R.M. and Moran,J.V. (2010) LINE-1 retrotransposition activity in human genomes. Cell, 141, 1159–1170.

7. Chuong,E.B., Elde,N.C. and Feschotte,C. (2017) Regulatory activities of transposable elements: from conflicts to benefits. Nat Rev Genet, 18, 71–86.

8. Attig,J., Agostini,F., Gooding,C., Chakrabarti,A.M., Singh,A., Haberman,N., Zagalak,J.A., Emmett,W., Smith,C.W.J., Luscombe,N.M., et al. (2018) Heteromeric RNP Assembly at LINEs Controls Lineage-Specific RNA Processing. Cell, 174, 1067–1081.e17.

9. Faulkner,G.J., Kimura,Y., Daub,C.O., Wani,S., Plessy,C., Irvine,K.M., Schroder,K., Cloonan,N., Steptoe,A.L., Lassmann,T., et al. (2009) The regulated retrotransposon transcriptome of mammalian cells. Nat Genet, 41, 563–571.

10. Marasca,F., Sinha,S., Vadalà,R., Polimeni,B., Ranzani,V., Paraboschi,E.M., Burattin,F.V., Ghilotti,M., Crosti,M., Negri,M.L., et al. (2022) LINE1 are spliced in non-canonical transcript variants to regulate T cell quiescence and exhaustion. Nat Genet, 54, 180–193.

11. Percharde,M., Lin,C.-J., Yin,Y., Guan,J., Peixoto,G.A., Bulut-Karslioglu,A., Biechele,S., Huang,B., Shen,X. and Ramalho-Santos,M. (2018) A LINE1-Nucleolin Partnership Regulates Early Development and ESC Identity. Cell, 174, 391–405.e19.

12. Treangen,T.J. and Salzberg,S.L. (2011) Repetitive DNA and next-generation sequencing: computational challenges and solutions. Nat Rev Genet, 13, 36–46.

13. Sexton,C.E. and Han,M.V. (2019) Paired-end mappability of transposable elements in the human genome. Mobile DNA, 10, 29.

14. Marasca,F., Gasparotto,E., Polimeni,B., Vadalà,R., Ranzani,V. and Bodega,B. (2020) The Sophisticated Transcriptional Response Governed by Transposable Elements in Human Health and Disease. International Journal of Molecular Sciences, 21, 3201.

15. Goerner-Potvin,P. and Bourque,G. (2018) Computational tools to unmask transposable elements. Nat Rev Genet, 19, 688–704.

16. Svensson,V., da Veiga Beltrame,E. and Pachter,L. (2020) A curated database reveals trends in single-cell transcriptomics. Database, 2020, baaa073.

17. He,J., Babarinde,I.A., Sun,L., Xu,S., Chen,R., Shi,J., Wei,Y., Li,Y., Ma,G., Zhuang,Q., et al. (2021) Identifying transposable element expression dynamics and heterogeneity during development at the single-cell level with a processing pipeline scTE. Nat Commun, 12, 1456.

18. Rodríguez-Quiroz,R. and Valdebenito-Maturana,B. (2022) SoloTE for improved analysis of transposable elements in single-cell RNA-Seq data using locus-specific expression. Commun Biol, 5, 1063.

19. Smith,T., Heger,A. and Sudbery,I. (2017) UMI-tools: modeling sequencing errors in Unique Molecular Identifiers to improve quantification accuracy. Genome Res., 27, 491–499.

20. Patro,R., Duggal,G., Love,M.I., Irizarry,R.A. and Kingsford,C. (2017) Salmon provides fast and bias-aware quantification of transcript expression. Nat Methods, 14, 417–419.

21. Bray,N.L., Pimentel,H., Melsted,P. and Pachter,L. (2016) Near-optimal probabilistic RNA-seq quantification. Nat Biotechnol, 34, 525–527.

22. Zhu,X., Fang,H., Gladysz,K., Barbour,J.A. and Wong,J.W.H. (2021) Overexpression of transposable elements is associated with immune evasion and poor outcome in colorectal cancer. European Journal of Cancer, 157, 94–107.

23. Kong,Y., Rose,C.M., Cass,A.A., Williams,A.G., Darwish,M., Lianoglou,S., Haverty,P.M., Tong,A.-J., Blanchette,C., Albert,M.L., et al. (2019) Transposable element expression in tumors is associated with immune infiltration and increased antigenicity. Nat Commun, 10, 5228.

24. Li,H., Handsaker,B., Wysoker,A., Fennell,T., Ruan,J., Homer,N., Marth,G., Abecasis,G., Durbin,R., and 1000 Genome Project Data Processing Subgroup (2009) The Sequence Alignment/Map format and SAMtools. Bioinformatics, 25, 2078–2079.

25. Kent,W.J., Sugnet,C.W., Furey,T.S., Roskin,K.M., Pringle,T.H., Zahler,A.M. and Haussler, and D. (2002) The Human Genome Browser at UCSC. Genome Res., 12, 996–1006.

26. Smit,A., Hubley,R. and Green,P. (2013) RepeatMasker Open-4.0.

27. Zheng,G.X.Y., Terry,J.M., Belgrader,P., Ryvkin,P., Bent,Z.W., Wilson,R., Ziraldo,S.B., Wheeler,T.D., McDermott,G.P., Zhu,J., et al. (2017) Massively parallel digital transcriptional profiling of single cells. Nat Commun, 8, 14049.

28. Kaminow,B., Yunusov,D. and Dobin,A. (2021) STARsolo: accurate, fast and versatile mapping/quantification of single-cell and single-nucleus RNA-seq data. biorXiv doi: https://doi.org/10.1101/2021.05.05.442755, 5 May 2021, pre-print: not peer-reviewed.

29. Dobin,A., Davis,C.A., Schlesinger,F., Drenkow,J., Zaleski,C., Jha,S., Batut,P., Chaisson,M. and Gingeras,T.R. (2013) STAR: ultrafast universal RNA-seq aligner. Bioinformatics, 29, 15–21.

30. Quinlan,A.R. and Hall,I.M. (2010) BEDTools: a flexible suite of utilities for comparing genomic features. Bioinformatics, 26, 841–842.

31. Frankish,A., Diekhans,M., Jungreis,I., Lagarde,J., Loveland,J.E., Mudge,J.M., Sisu,C., Wright,J.C., Armstrong,J., Barnes,I., et al. (2021) GENCODE 2021. Nucleic Acids Res, 49, D916–D923.

32. 10x Genomics (2017) 8k PBMCs from a Healthy Donor, Single Cell Gene Expression Dataset by Cell Ranger 2.1.0.

33. Li,H. (2013) Aligning sequence reads, clone sequences and assembly contigs with BWA-MEM. biorXiv doi: https://doi.org/10.48550/arXiv.1303.3997, 26 May 2013, pre-print: nor peer-reviewed.

34. R core Team (2021) R: A Language and Environment for Statistical Computing.

35. Hao,Y., Hao,S., Andersen-Nissen,E., Mauck,W.M., Zheng,S., Butler,A., Lee,M.J., Wilk,A.J., Darby,C., Zager,M., et al. (2021) Integrated analysis of multimodal single-cell data. Cell, 184, 3573–3587.e29.

36. Wei,T. and Simko,V. (2021) R package ‘corrplot’: Visualization of a Correlation Matrix.

37. Di Tommaso,P., Chatzou,M., Floden,E.W., Barja,P.P., Palumbo,E. and Notredame,C. (2017) Nextflow enables reproducible computational workflows. Nat Biotechnol, 35, 316–319.

38. Lee,H.-O., Hong,Y., Etlioglu,H.E., Cho,Y.B., Pomella,V., Van den Bosch,B., Vanhecke,J., Verbandt,S., Hong,H., Min,J.-W., et al. (2020) Lineage-dependent gene expression programs influence the immune landscape of colorectal cancer. Nat Genet, 52, 594–603.

39. Robinson,J.T., Thorvaldsdóttir,H., Winckler,W., Guttman,M., Lander,E.S., Getz,G. and Mesirov,J.P. (2011) Integrative genomics viewer. Nat Biotechnol, 29, 24–26.

40. Lanciano,S. and Cristofari,G. (2020) Measuring and interpreting transposable element expression. Nat Rev Genet, 21, 721–736.

41. Mortazavi,A., Williams,B.A., McCue,K., Schaeffer,L. and Wold,B. (2008) Mapping and quantifying mammalian transcriptomes by RNA-Seq. Nat Methods, 5, 621–628.

42. Wolf,F.A., Angerer,P. and Theis,F.J. (2018) SCANPY: large-scale single-cell gene expression data analysis. Genome Biology, 19, 15.

43. Srivastava,A., Malik,L., Smith,T., Sudbery,I. and Patro,R. (2019) Alevin efficiently estimates accurate gene abundances from dscRNA-seq data. Genome Biology, 20, 65.

44. Melsted,P., Booeshaghi,A.S., Liu,L., Gao,F., Lu,L., Min,K.H. (Joseph), da Veiga Beltrame,E., Hjörleifsson,K.E., Gehring,J. and Pachter,L. (2021) Modular, efficient and constant-memory single-cell RNA-seq preprocessing. Nat Biotechnol, 39, 813–818.

45. Lun,A.T.L., Riesenfeld,S., Andrews,T., Dao,T.P., Gomes,T., Marioni,J.C., and participants in the 1st Human Cell Atlas Jamboree (2019) EmptyDrops: distinguishing cells from empty droplets in droplet-based single-cell RNA sequencing data. Genome Biology, 20, 63.

46. Harris,C.R., Millman,K.J., van der Walt,S.J., Gommers,R., Virtanen,P., Cournapeau,D., Wieser,E., Taylor,J., Berg,S., Smith,N.J., et al. (2020) Array programming with NumPy. Nature, 585, 357–362.

47. Bonfield,J.K., Marshall,J., Danecek,P., Li,H., Ohan,V., Whitwham,A., Keane,T. and Davies,R.M. (2021) HTSlib: C library for reading/writing high-throughput sequencing data. GigaScience, 10, giab007.

48. Heger,A. and Jacobs,K. (2020) Pysam.

49. Satija,R., Farrell,J.A., Gennert,D., Schier,A.F. and Regev,A. (2015) Spatial reconstruction of single-cell gene expression data. Nat Biotechnol, 33, 495–502.

50. Jang,H.S., Shah,N.M., Du,A.Y., Dailey,Z.Z., Pehrsson,E.C., Godoy,P.M., Zhang,D., Li,D., Xing,X., Kim,S., et al. (2019) Transposable elements drive widespread expression of oncogenes in human cancers. Nat Genet, 51, 611–617.

51. Kines,K.J., Sokolowski,M., deHaro,D.L., Christian,C.M. and Belancio,V.P. (2014) Potential for genomic instability associated with retrotranspositionally-incompetent L1 loci. Nucleic Acids Res, 42, 10488–10502.

52. Goodier,J.L. (2016) Restricting retrotransposons: a review. Mobile DNA, 7, 16.

53. Nicolae,M., Mangul,S., Măndoiu,I.I. and Zelikovsky,A. (2011) Estimation of alternative splicing isoform frequencies from RNA-Seq data. Algorithms for Molecular Biology, 6, 9.

54. Li,W., Prazak,L., Chatterjee,N., Grüninger,S., Krug,L., Theodorou,D. and Dubnau,J. (2013) Activation of transposable elements during aging and neuronal decline in Drosophila. Nat Neurosci, 16, 529–531.

55. Burns,K.H. (2017) Transposable elements in cancer. Nat Rev Cancer, 17, 415–424.

56. Ochoa Thomas,E., Zuniga,G., Sun,W. and Frost,B. (2020) Awakening the dark side: retrotransposon activation in neurodegenerative disorders. Current Opinion in Neurobiology, 61, 65–72.

